# A conservative approach for species delimitation based on multi-locus DNA sequences: a case study of the genus *Giraffa* (Mammalia, Cetartiodactyla)

**DOI:** 10.1101/648162

**Authors:** Alice Petzold, Alexandre Hassanin

## Abstract

Molecular data are now commonly used in taxonomy for delimiting cryptic species. In the case of giraffes, which were treated as a single species (*Giraffa camelopardalis*) during half of a century, several molecular studies have suggested a splitting into four to seven species, but the criteria applied for taxonomic delimitation were not fully described.

In this study, we have analysed all multi-locus DNA sequences available for giraffes using multispecies coalescent (MSC: *BEAST, BPP and STACEY), population genetic (STRUCTURE, allelic networks, haplotype network and bootstrapping) and phylogenetic (MrBayes, PhyML, SuperTRI) methods to identify the number of species. Our results show that depending on the method chosen, different taxonomic hypotheses, recognizing from two to six species, can be considered for the genus *Giraffa.* Our results confirm that MSC methods can lead to taxonomic over-splitting, as they delimit geographic structure rather than species. The 3-species hypothesis, which recognizes *G. camelopardalis* sensu strico, *G. giraffa*, and *G. tippelskirchi*, is highly supported by phylogenetic analyses and also corroborated by most population genetic and MSC analyses. The three species show high levels of nucleotide divergence in both nuclear (0.35-0.51 %) and mitochondrial sequences (3-4 %), and they are characterised by 7 to 12 exclusive synapomorphies (ES) detected in nine of the 21 nuclear introns analysed for this study. By contrast, other putative species, such as *G. peralta*, *G*. *reticulata*, *G. thornicrofti* or *G. tippelskirchi* sensu stricto, do not exhibit any ES in nuclear genes.

A robust mito-nuclear conflict was found for the position and monophyly of *G. giraffa* and *G. tippelskirchi*, which is explained firstly by a mitochondrial introgression from Masai giraffe to southeastern giraffe during the Pleistocene, and secondly, by gene flow mediated by male dispersal between southern populations (subspecies *G.g. giraffa* and *G.g. angolensis*).

## Introduction

Biologically, speciation implies reproductive isolation through barriers preventing or limiting gene flow between populations [1]. Over the process of genetic differentiation, reproductively isolated populations may accumulate distinct phenotypic features that facilitate their recognition as different species. However, separated populations facing similar selective environments often converge phenotypically and show no visible differences (see Fišer et al. [2] for a review on cryptic species).

For more than three decades, mitochondrial genes, and in particular the *COX1* gene (cytochrome c oxidase subunit 1), have been intensively used for species delimitation [3–4]. However, numerous molecular studies have revealed that the mitochondrial tree may deviate from the species tree. Indeed, the maternal inheritance of the mtDNA genome can be misleading for species delimitation because females and males have generally different dispersal behaviours (female philopatry versus male dispersal) [5,6], and because interspecific hybrid females are generally fertile, whereas hybrid males are often sterile (Haldane’s rule), facilitating mitochondrial introgression between closely related species [7–9]. To overcome these limitations, most recent taxonomic studies dealing with the delimitation between cryptic mammal species have focused on multi-locus datasets [10–12], as the use of multiple independent DNA markers has been shown to provide a strong and reliable signal for deciphering relationships among closely related taxa [13–14]. However, interpreting the results from multi-locus datasets can be difficult, especially when the DNA markers show low genetic variation or conflicting relationships between them. These difficulties have led to the development of a plethora of new methodological approaches for multi-locus species delimitation [15,16], which may be subdivided into three categories: (1) phylogenetic methods, (2) multispecies coalescent (MSC) approaches, and (3) population genetic methods (Table 1). Phylogenetic methods were not originally developed for studying species delimitation, but the species monophyly criterion has been widely used since the origin of molecular taxonomy [17]. For multi-locus datasets, several phylogenetic approaches can be considered: the concatenation of all markers into a supermatrix (although this approach has been widely criticized [18]), the separate analyses of the markers, or more sophisticated methods, such as *BEAST [19] or SuperTRI [20]. Based on the coalescent theory, some authors have suggested that species can be delimited without monophyletic gene trees [21].

**Table 1.** Species delimitation methods based on multi-locus nuDNA sequences used in this study

| Method | Input | Category | Reference | Description | SD Criteria |
| --- | --- | --- | --- | --- | --- |
| **STRUCTURE** | alignment of PHASED alleles | PG | Pritchard et al. [26] ;<br>Falush et al. [46] | Bayesian clustering method based on the estimation of allele frequencies | $\Delta K$ and plateau methods |
| **BPP**<br>(Bayesian Phylogenetics and Phylogeography) | alignment of consensus sequences | MCS | Yang and Rannala [55] | Bayesian method based on the MSC model, in which a reversible-jump Markov chain Monte Carlo algorithm is used to calculate the posterior probabilities of species delimitations. | Probability $\geq 0.95$ |
| **STACEY**<br>(Species Tree and Classification Estimation, Yarely) | alignment of PHASED alleles | MCS | Jones [24]<br>-----<br>Individual assignment of alleles: present study | Bayesian method implemented in BEAST 2 [56] for the inference of a “species or minimal clusters tree” (SMC) under the birth-death-collapse tree prior and without the requirement of a guide tree. | Probability $\geq 0.95$ |
| **\*BEAST**<br>(Species Tree Ancestral Reconstruction in BEAST) | alignment of PHASED alleles | MCS<br>P | Program: Heled and Drummond [19];<br>Individual assignment of alleles: present study | Bayesian method implemented in BEAST 2 [56] based on the MSC model | Probability $\geq 0.95$ |
| **Bootstrap Analysis of Haplotypes** | alignment of PHASED haplotypes | P<br>PG | Present study | Bootstrap consensus tree reconstructed with ML, MP or NJ methods | Bootstrap $\geq 90$ |
| **Supermatrix**<br>MrBayes | alignment of consensus sequences | P | Ronquist et al. [40] | Bayesian inference of phylogeny | Probability $\geq 0.95$ |
| PhyML | | | Guindon et al. [41] | ML method for tree construction | Bootstrap $\geq 90$ |
| **SuperTRI**<br>(SuperTree with Reliability Indices) | Weighted binary matrix of node support for each locus | P | Ropiquet et al. [20] | Three measures are calculated to estimate the reliability of the nodes (SBP, MPP and NRep) using the branch support values (PP) of all phylogenetic hypotheses produced during the separate Bayesian analyses of the 21 introns | (1) SBP $\geq 90$<br>(2) MPP $\geq 0.1$<br>(3) NRep $\geq 2$ |

SD: Species delimitation; PG: Population Genetics; MSC: Multispecies Coalescent; P: Phylogenetic methods; SBP: Supertree Bootstrap Percentage; MPP: Mean Posterior Probability; Rep: Reproducibility Index; SPR: Subtree Pruning and Re-grafting

The incorporation of the coalescent model [22] in certain software (e.g. *BEAST [19], BPP [23] and STACEY [24]) enabled the inference of species limits from multi-locus data by accounting for incongruences among gene trees in the presence of incomplete lineage sorting [19]. MSC approaches often require prior assignments of samples to populations or taxa and are hence restricted to the validation of proposed delimitations [25]. Population genetic approaches are generally applied to detect “cryptic substructure” between groups showing very similar phenotypes. The program STRUCTURE [26] is probably the most popular approach for Bayesian clustering using multi-locus data. It has recently gained new interest as the clusters identified with STRUCTURE can be used as preliminary hypothesis for assigning individuals to populations or taxa, which represents the first step of most MSC analyses [27]. In addition, geographic clusters detected with STRUCTURE are often interpreted (perhaps wrongly [28]), as reproductively isolated populations, which may constitute a strong argument in favour of a division into several species (e.g., Brown et al. [29]).

The systematics of giraffes is a controversial issue, since at least nine different hypotheses of species delimitation were proposed on the basis of morphological characters and, more recently, molecular data (Appendix A1). The existence of several giraffe species was first proposed by Geoffroy Saint-Hilaire [30], who noted that differences in coat pattern, horn shape and skull can be used to distinguish the Nubian giraffe (from the Sennaar region in Sudan) from the Southern giraffe (from the Cape region). Thomas [31] proposed another arrangement in two species, in which Nubian and Southern giraffes were assigned to *Giraffa camelopardalis*, whereas the reticulated giraffe was treated as a full species, *Giraffa reticulata*. Lydekker [32] shared this view, but recognized 12 subspecies in *G. camelopardalis* and two in *G. reticulata*. However, Dagg and Foster [33] indicated that phenotypic features are highly variable between and within populations, and recognized therefore a single species, *camelopardalis*. Subsequently, this point of view was accepted by most other taxonomists, despite persisting controversy regarding the number of subspecies [34,37]. However, the taxonomy of giraffes has been challenged by recent genetic studies: based on the analyses of mitochondrial sequences and 14 nuclear microsatellite loci, Brown et al. [29] proposed a minimum of six species, corresponding to *Giraffa angolensis*, *G. giraffa*, *G. peralta*, *G. reticulata*, *G. rothschildi*, and *G. tippelskirchi* (N.B. the subspecies *camelopardalis*, *antiquorum* and *thornicrofti* were not included in their study); whereas Fennessy et al. [38] and Winter et al. [12] suggested a division into four species, i.e., *G. camelopardalis*, *G. giraffa*, *G. reticulata* and *G. tippelskirchi*, based on multi-locus analyses of 7 and 21 nuclear introns, respectively.

In this study, we reanalysed all multi-locus data available for the nine giraffe subspecies (i.e., *camelopardalis*, *angolensis*, *antiquorum*, *giraffa*, *peralta*, *reticulata*, *rothschildi*, *thornicrofti* and *tippelskirchi*; see geographic distributions in Fig. 2) using various phylogenetic (MrBayes, PhyML, SuperTRI), population genetic (STRUCTURE, allelic networks, haplotype network and bootstrapping) and MSC (*BEAST, BPP and STACEY) methods. Our five main goals were (1) to test if the different methods converge towards the same conclusion or if they support divergent taxonomic hypotheses, (2) to examine if one hypothesis is more supported by the analyses than the others (conservative approach of species delimitation), (3) to understand why some methods or models can lead to taxonomic over-splitting, (4) to know if available molecular data are sufficient to conclude on the number of species, and (5) to determine which data, methods and operational criteria are relevant for delimiting species with molecular data.

**Figure 1.**
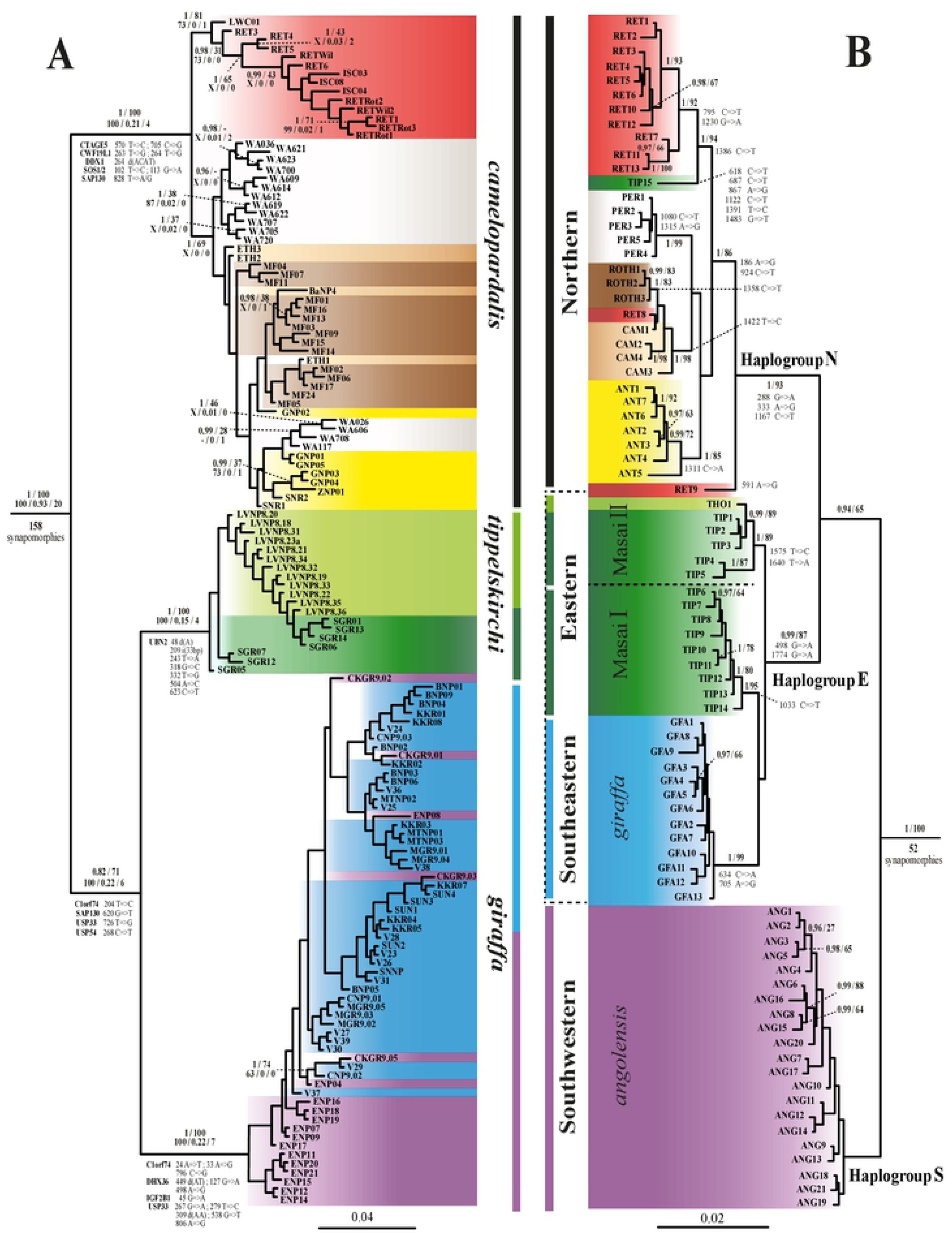
Comparative phylogeny of nuclear and mitochondrial datasets. The nine subspecies are differentiated by the following colours: red: *reticulata*, white: *peralta*, brown: *rothschildi*, beige: *camelopardalis*, yellow: *antiquorum*, blue: *giraffa*, purple: *angolensis*, light green: *thornicrofti* and dark green: *tippelskirchi*. The three outgroup species are not shown. (A) Bayesian tree inferred from the nuclear dataset, named nuDNA-G137O3, including the sequences of 21 introns for 137 giraffes. The tree was rooted with *Bos*, *Ovis*, and *Okapia* (not shown). For each node recovered with significant support in the Bayesian analysis (PP ≥ 0.9), as well as for other nodes discussed in the text, the two values above indicate the Posterior Probability with MrBayes (PP) and the Bootstrap Percentage obtained from the Maximum Likelihood analysis (BP). The three values below were obtained from the SuperTRI analyses of the 21 introns: from left to right: Supertree Bootstrap Percentage (SBP), Mean Posterior Probability (MPP) and the number of markers supporting the node (NRep). The symbol ‘‘–’’ indicates that the node was not found monophyletic in the analysis, and the letter ‘‘X’’ indicates that an alternative hypothesis was supported by SBP > 50. The exclusive synapomorphies (including indels; i: insertion; d: deletion), representing fixed substitutions among members of a group, are listed for the nodes discussed in the text. (B) Bayesian tree of the 82 mitochondrial haplotypes detected for *Giraffa* reconstructed from a fragment covering the complete *Cytb* gene and the 5’ part of the control region (1776 characters) and rooted with *Bos*, *Ovis* and *Okapia* (not shown). For each node supported by PP ≥ 0.95, the BP value obtained from the Maximum Likelihood analysis is indicated. Fixed substitutions among members of a group (exclusive synapomorphies) are listed for the nodes supported by PP ≥ 0.95 and for uncommon mitochondrial haplotypes.

**Figure 2.**
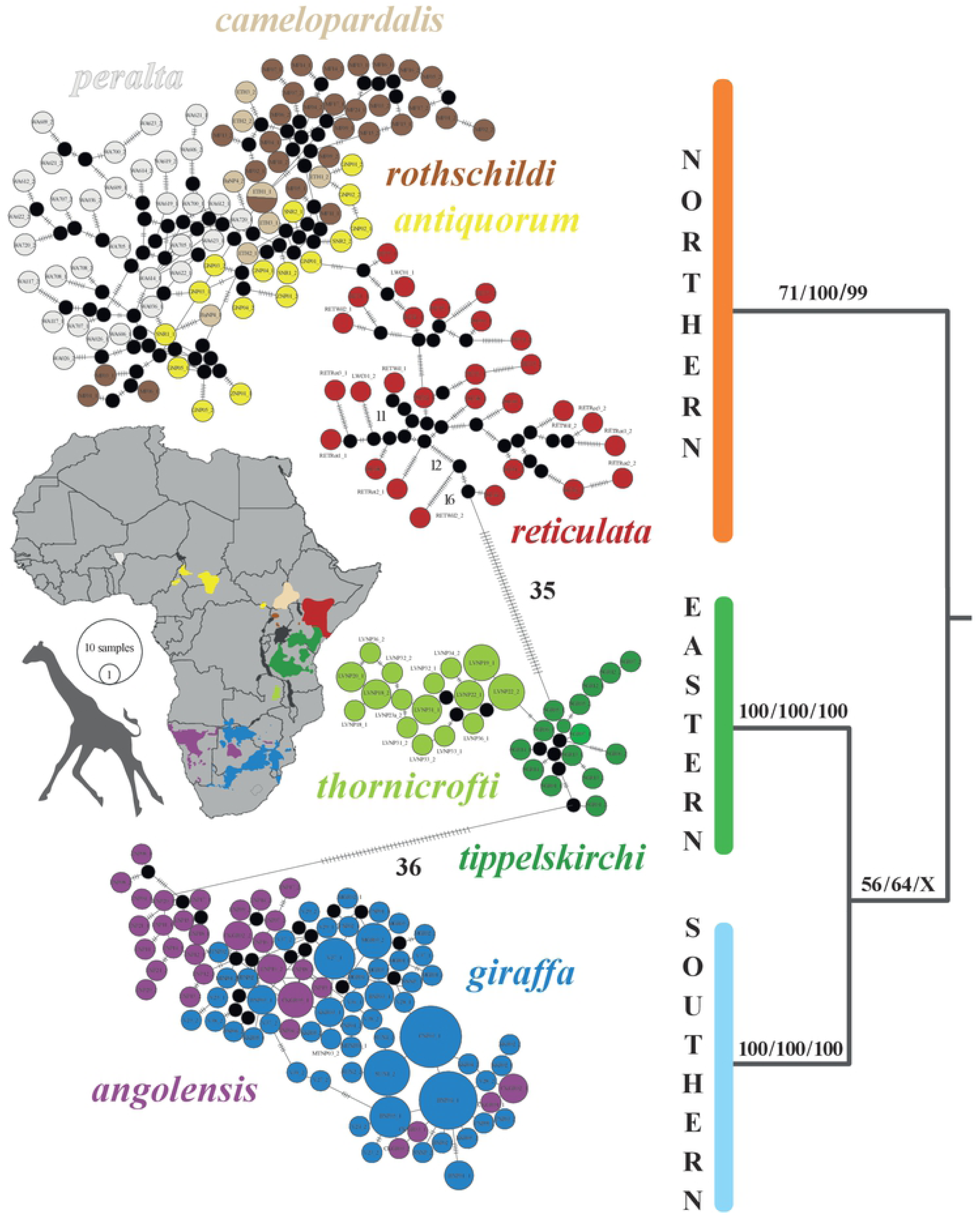
Current distribution of giraffe subspecies and population genetic analyses of nuclear haplotypes. The nine subspecies currently recognized are distinguished by different colours on the map (modified from https://giraffeconservation.org/giraffe-species/). At the left, the median-joining network was constructed under PopART using the nuDNA-G274 dataset, which corresponds to the 274 nuclear haplotypes inferred under PHASE for the 137 giraffes sequenced for 21 introns. The numbers of mutations between haplotypes are indicated on the branches. At the right, the 50% majority-rule bootstrap consensus tree was reconstructed under PAUP using the nuDNA-G274O6 dataset (see Material and Methods for more details). The values at the nodes represent Bootstrap percentages ≥ 50 calculated with maximum parsimony, distance and maximum likelihood methods (from left to right). Relationships within subspecies are not shown here.

## Material and Methods

### Nuclear and mitochondrial datasets used for the analyses

Seven giraffe datasets were generated for our analyses using the sequences available in the NCBI nucleotide database:

1. the mtDNA-G507 dataset, which contains a mitochondrial fragment covering the whole cytochrome b (*Cytb*) gene and the 5‘part of the control region (length = 1742 nucleotides [nt]) for 507 individuals (listed in Appendix B1.1), and its reduced version including only the 82 different mitochondrial haplotypes, named mtDNA-GH82;
2. the mtDNA-GH82O3, in which the mtDNA-GH82 dataset was aligned to three outgroup species: *Bos taurus* (NCBI accession number KT184464); *Ovis canadensis* (NC_015889) and *Okapia johnstoni* (JN632674) (length = 1776 nt);
3. a nuclear dataset, named nuDNA-G274, including 274 phased alleles of 21 introns (ACP5, C1orf74, CCT2, COL5A2, CTAGE5, CWF19L1; DDX1, DHX36, IGF2B1, MACF1, NOTCH2, NUP155, OTOF, PLCE1, RASSF4, RFRC5, SAP130, SOS1, UBN2, USP33, USP54) for 137 giraffes (accession numbers LT596685-LT598170, MG257969–MG262280);
4. the nuDNA-G274O6, in which the nuDNA-G274 dataset was aligned to the alleles of three outgroup species: the okapi (*Okapia johnstoni*, published sequences [12,38]), and two bovid species, (i) *Bos taurus*, for which the sequences were extracted by BLAST from the whole genome version UMD3.1.1 (http://bovinegenome.org/) or, in case of unavailability of certain genes, from the genome of *Bos mutus* available on NCBI (SAMN08580377); and (ii) *Ovis canadensis*, for which the sequences were extracted by BLAST from the genome available on NCBI (CP011888.1);
5. the nuDNA-G137 dataset, comprising the alignments of original consensus sequences of 21 introns for the 137 giraffes (length = 16968 nt), which were recovered by detecting heterozygous sites in Geneious R10 (Biomatters, Auckland, New Zealand);
6. the nuDNA-G137O3 dataset, in which the nuDNA-G137 dataset was aligned to the three outgroup species mentioned above (length = 17276 nt);
7. the nuclear haplotype dataset, named nuDNA-GH274, which was inferred from the nuDNA-G274 dataset using the PHASE v2.1 algorithm implemented in the software DNASP v5.0 [39] (length = 1362 nt; it contains only the sites found to be variable between giraffe haplotypes).

All alignments generated for this study were deposited in DRYAD (entry doi: XXXXXXX).

### Phylogenetic analyses

The mtDNA-GH82O3 and nuDNA-G137O3 datasets were analysed with probabilistic methods. Bayesian inferences were conducted in MrBayes v3.2.6 [40] by calculating the posterior probabilities (PP) after 10^7^ Metropolis-coupled MCMC generations with tree sampling every 1000 generations and a burn-in of 25 %. Maximum Likelihood (ML) analyses were performed with PhyML v3.1 [41] and Bootstrap percentages (BP) were calculated after 1000 replicates. The GTR+I+G substitution model was applied for both methods, as suggested by the Likelihood calculations in jModeltest [42] based on the Akaike information criterion.

Bayesian analyses were also performed for each of the 21 introns using the model of DNA substitution selected under jModeltest (Table 1).

### SuperTRI analyses

The lists of bipartitions obtained from the Bayesian analyses (.parts and .tstat files) for each nuclear marker were transformed into a weighted binary matrix (MRP, matrix representation with parsimony) for supertree construction using SuperTRI v57 [20]. Here, each binary character corresponds to a node, which was weighted according to its frequency in one of the 21 lists of bipartition. Thereby, the SuperTRI method accounts for principal as well as secondary signals, given that all phylogenetic hypotheses found during the Bayesian analyses are represented in the weighted binary matrix used for supertree construction. The reliability of the nodes was assessed using three measures: supertree bootstrap percentages (SBPs) were obtained from PAUP* v4b10 [43] after 1000 BP replicates of the MRP matrix of 24749 binary characters generated by SuperTRI v57; mean posterior probabilities (MPP) and reproducibility indices (Rep) were directly calculated on SuperTRI v57. In the nuclear tree (Fig. 1A), we chose to indicate the number of markers supporting each node of interest (NRep) rather than the Rep value, which represents the ratio of the number of markers supporting the node to the total number of markers [20].

### STRUCTURE analyses

Giraffe haplotypes were reconstructed from the nuDNA-G137O3 dataset for each of the 21 introns by applying the PHASE v2.1 algorithm implemented in the software DNASP v5.0 [39], allowing for recombination and reducing the output probability threshold of conserved regions (CT) from 0.9 by default to 0.6. For each of the 21 introns, the haplotype information was used to code individuals sharing the same allele with a unique integer.

Bayesian analyses of genetic admixture were run in STRUCTURE v.2.3.4 [44] to identify genetically homogeneous groups of individuals (populations of origin, K). The analyses were done as recommended by Gilbert et al. [45], i.e., number of MCMC generations = 200 000 and burn-in = 100 000 generations for K = 1-10 clusters. We applied several combinations of ancestry model, allele frequency and supporting information (Popdata) like the assignment of the subspecies (population identity/ POPID) or sampling location (LOCPRIOR model) for each individual. We tested two ancestry models, since we do not know whether studied populations were discrete or had an admixed ancestry. Moreover, the identification of the most probable number of clusters (K) might be further affected by the choice of the allele frequency model. By default, the software assumes correlated allele frequency among populations caused by migration and shared ancestry [46]. Since past admixture was expected between giraffe populations, this model may represent the appropriate choice. However, several runs were conducted under the independent allele frequency model, as it might be more powerful to detect highly distinct populations [47].

We also tested two settings for lambda (λ), the parameter specifying the distribution of allelic frequencies in each population: the default setting (λ = 1) and an estimated value of λ (λ = 0.45), calculated during a run comprising 20 iterations for K = 1. Runs were performed without any assignation of individuals, or by assigning individuals to either a POPID representing the designated subspecies or to their sampling location (LOCPRIOR, national parks where the giraffes were sampled; Appendix B2), as this option is recommended when only a weak signal is present in the markers [48]. All analyses were replicated 20 times.

The most likely number of distinct groups for each run was identified by means of STRUCTURE HARVESTER [49]. Thereby, the optimal K was determined using two approaches: (1) the ΔK method of Evanno et al. [50], which recognizes the most likely number of distinct clusters by the largest ΔK value, calculated by the rate of change in the log probability of data between successive K values; and (2) the “plateau “ method of Pritchard et al. [44], where the log probability of the data (ln Pr (X|K) was plotted against a range of K values, and the optimal K was selected as the point at which the plot curvature plateaus. A regression curve and gridlines were added to the diagrams generated by STRUCTURE HARVESTER to help in determining the point of plateau.

To assess the reliability of the results, CLUMPAK [51] was used to display the barplots from K = 1 to 10 for each of the 20 iterations by means of the implemented software DISTRUCT [52].

### Analyses of nuclear haplotypes

The nuDNA-GH274 dataset (nuclear haplotypes inferred for 21 introns and 137 giraffes) was used to construct a median joined network using PopART v1.7 following the distance criterion [53]. The robustness of haplotype clusters was evaluated by bootstrapping (1000 replicates) under PAUP* v4b10 [43] using either the Maximum Parsimony (MP) method (heuristic search, faststep option) or the Neighbor-Joining (NJ) method (GTR+I+G model), and under the Maximum Likelihood (ML) criterion using RAxML on CIPRES [54] (http://www.phylo.org).

PopART was also used to construct a median joined network for each of the 21 introns. For six introns (i.e. *CTAGE5*, *NUP155, OTOF, PLCE1, RASSF4* and *SOS1*), missing alleles were removed from the alignment to avoid any distortion of the results.

### Multispecies coalescent analyses

Three coalescent-based approaches were applied to infer species boundaries within the genus *Giraffa*: (1) the “Species Tree Ancestral Reconstruction” template (*BEAST [19]), (2) the extension of the *BEAST model called “Species Tree and Classification Estimation, Yarely” (STACEY) [24], and (3) the Bayesian Phylogenetics and Phylogeography program (BPP v.3.2 [23,55]) (see specifications for each program in Table 1).

We estimated the species-tree phylogeny using the coalescent algorithm implemented in BEAST v.2.4.4 [56] in order to consider an alternative to the traditional concatenated phylogenetic approach (see Kubatko and Degnan [57] for caveats concerning concatenation). Inferences were based on the nuDNA-G274O6 dataset using an *a priori* assignment at the level of individuals, i.e. by assigning for each of the 137 giraffes two alleles for the 21 introns. We assumed an uncorrelated lognormal molecular clock for all 21 loci. For each marker, we selected the best suited substitution model inferred in jModeltest [42] (Table 2). Analyses were run with 2x 10^8^ generations, with trees sampled every 5000 steps. The .log files were analysed with Tracer v1.7 [58] to assess the convergence of model parameters (effective sample size [ESS] > 200). The species tree was summarized as a Maximum Clade Credibility tree in TreeAnnotator v.1.10 [59] after discarding 25% as burn-in.

**Table 2.** Characteristics of the nuclear alignments used for Bayesian phylogenetic analyses and posterior probabilities obtained for the different giraffe taxa.

| Alignments | Substitution Model* | Length* | IS* | C | R | T | G | CR | CRG | CRT | GT | Th |
| --- | --- | --- | --- | --- | --- | --- | --- | --- | --- | --- | --- | --- |
| ACP5 | F81+G | 640 | 9 | - | 0.03 | - | 0.04 | 0.75 | - | 0.52 | - | - |
| C1orf74 | F81 | 870 | 9 | - | - | - | 1.00 | - | - | - | 1.00 | - |
| CCT2 | HKY | 812 | 6 | - | - | - | - | - | - | - | - | - |
| COL5A2 | HKY | 878 | 7 | - | - | 0.96 | - | - | - | - | 0.04 | - |
| CTAGE5 | HKY | 839 | 13 | - | - | 0.64 | 0.10 | 1.00 | - | - | - | - |
| CWF19L1 | HKY | 668 | 5 | - | - | - | - | 1.00 | - | - | - | - |
| DDX1 | HKY | 734 | 15 | - | - | - | - | - | - | - | 0.43 | - |
| DHX36 | HKY | 815 | 11 | - | - | - | 0.98 | - | - | 1.00 | - | - |
| IGF2B1 | HKY+G | 802 | 7 | - | - | - | 0.90 | - | - | - | - | 1.00 |
| MACF1 | HKY | 718 | 9 | - | - | - | - | - | - | - | - | - |
| NOTCH2 | F81 | 854 | 5 | - | - | - | 0.50 | - | - | - | - | - |
| NUP155 | HKY | 659 | 5 | - | - | - | - | - | - | - | - | - |
| OTOF | K80 | 741 | 7 | - | - | - | - | - | - | - | - | - |
| PLCE1 | HKY | 836 | 9 | - | - | - | - | - | - | - | - | - |
| RASSF4 | SYM+G | 646 | 10 | - | - | - | - | - | - | - | - | - |
| RFC5 | HKY+G | 825 | 9 | - | - | 0.75 | - | - | 0.70 | - | - | - |
| SAP130 | HKY+G | 888 | 11 | - | - | - | - | - | - | - | - | - |
| SOS1 | F81 | 758 | 4 | - | - | - | - | 1.00 | - | - | - | - |
| UBN2 | F81 | 719 | 11 | - | - | 1.00 | - | - | 1.00 | - | - | - |
| USP33 | HKY+G | 937 | 13 | - | - | - | 1.00 | - | - | - | 0.98 | - |
| USP54 | HKY | 1329 | 12 | - | - | - | - | - | - | - | 0.98 | - |
| nuDNA | GTR+I+G | 16968 | 187 | 1.00 | 1.00 | 1.00 | 1.00 | 1.00 | - | - | 0.82 | - |

IS: Informative sites for parsimony within *Giraffa*; C: *G. camelopardalis* sensu stricto A (see Fig. 4); R: *G. reticulata*; T: *G. tippelskirchi* (subspecies *tippelskirchi* and *thornicrofti*); G: *G. giraffa*; Th: *G. thornicrofti*; P: *G. peralta*; "-": not found; "\*": outgroups excluded.

The nuDNA-G274 and nuDNA-G274O6 datasets were further used for species delimitation analyses using the STACEY template implemented in BEAST v2.4.4. STACEY represents an improvement to the DISSECT model [60] to infer a “species or minimal clusters tree” (SMC) under the birth-death-collapse tree prior and without the requirement of a guide tree. The tips of the SMC tree represent minimal clusters of individuals that may be collapsed to a single putative species, if branches are shorter than a specified length (collapse height) [24]. A first run was conducted without taxonomic a priori assumptions by assigning two alleles per gene and individual. For the other run, each individual was assigned to one of the six taxa (6S hypothesis) that were found monophyletic with at least one of our phylogenetic analyses: *G. camelopardalis* sensu stricto C (including only the three subspecies *camelopardalis, antiquorum* and *rothschildi*), *G. peralta, G. reticulata, G. giraffa* (including the two subspecies *angolensis* and *giraffa*), *G. tippelskirchi* sensu stricto and *G. thornicrofti.* Analyses were done as suggested in the manual, i.e. using a relative death rate of 0.0 for the tree prior, a lognormal distribution with a mean of 4.6 and a standard deviation of 2 to the growth rate prior and a uniform distribution for the relative death rate prior with a lower bound of −0.5 and an upper bound of 0.5. The dataset was partitioned by the 21 genes, with independent strict clock models and individual assignment of the best suited substitution model to each gene (Table 2). Each analysis was run for 2.5 x 10^8^ generations and convergence of parameters was assessed in Tracer v1.7 [58]. Subsequently, the most supported number of distinct clusters was estimated using SpeciesDelimitationAnalyser v1.8.0 [24] by analysing the species trees with a burn-in of 25 % and the default collapse height of 0.0001.

Species delimitation analyses with BPP v3.2 were based on a reduced dataset comprising only 66 giraffes due to software limitation. Seventy-one individuals were excluded from the original dataset using the three following criteria: (1) 14 individuals with missing data, (2) 39 individuals sharing the same haplotype and (3) 18 individuals characterized by a long terminal branch in the Bayesian tree. We analysed the support for each of the five taxonomic hypotheses depicted in Fig. 4 (see results for more details). First, we applied the A00 algorithm, the simple MSC model with the species tree fixed to explain the acceptance proportions of MCMC moves [23] under the default gamma prior values *G* (2, 2000) for the tau (τ, root divergence time) and theta (θ, genetic difference among taxa). Then, we assessed the support for each putative species using the A11 algorithm [55]. The three species model priors (SMP 1, 2 and 3) were tested. The analyses were run for 500 000 generations followed by a burn-in of 10 %. Convergence between runs was checked for fine tune acceptance proportions between 0.15 and 0.7, as well as ESS *>*200.

**Figure 3.**
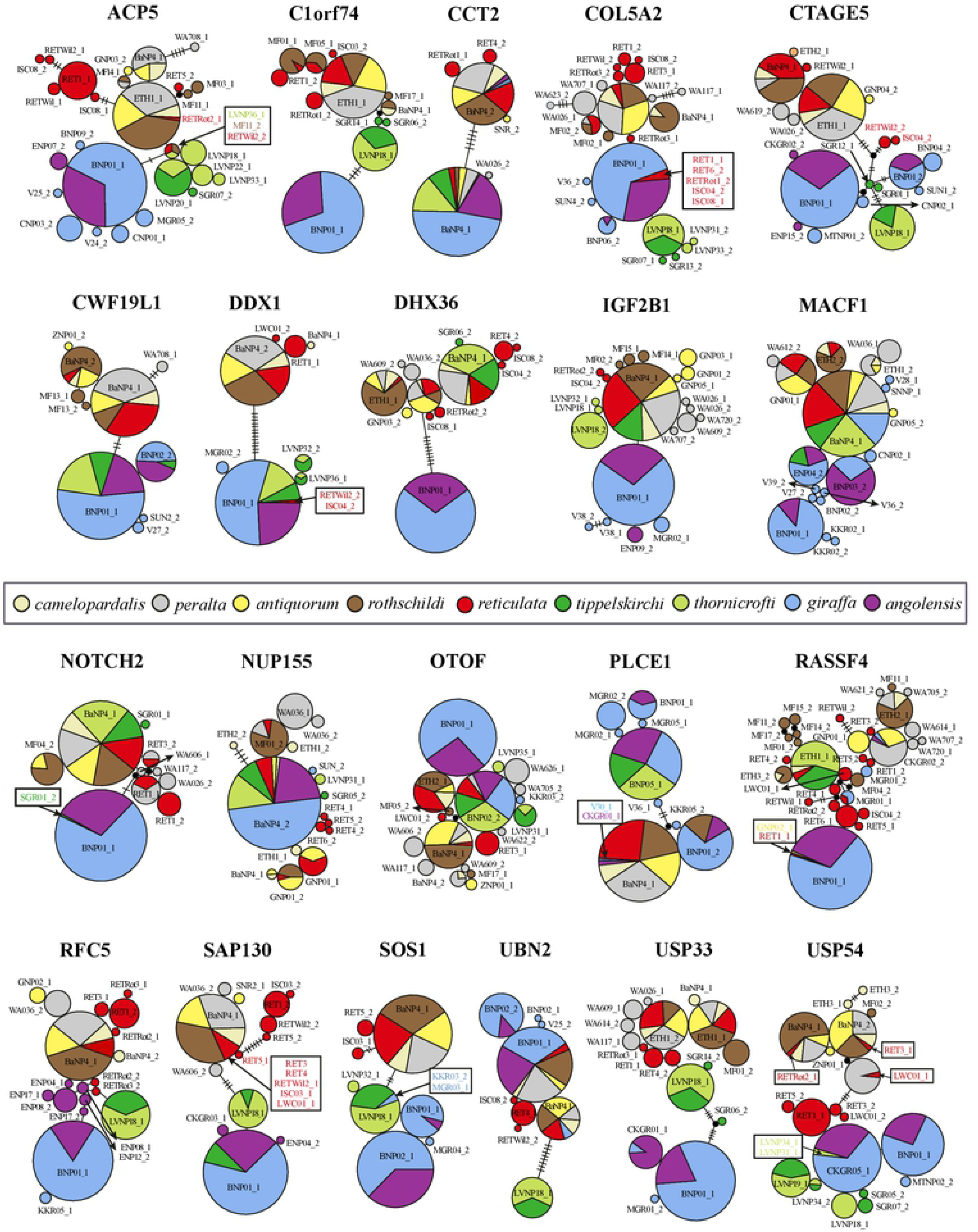
Allelic networks for 21 nuclear introns. The circles represent alleles with sizes proportional to their frequency in the populations. Each allele is designated with one representative individual (the list of all individuals is provided in Appendix F). The nine subspecies currently recognized are distinguished by different colours. Individuals characterized by a rare allele (in the subspecies) are highlighted with a black frame. The numbers of mutations between alleles are indicated on the branches.

**Figure 4.**
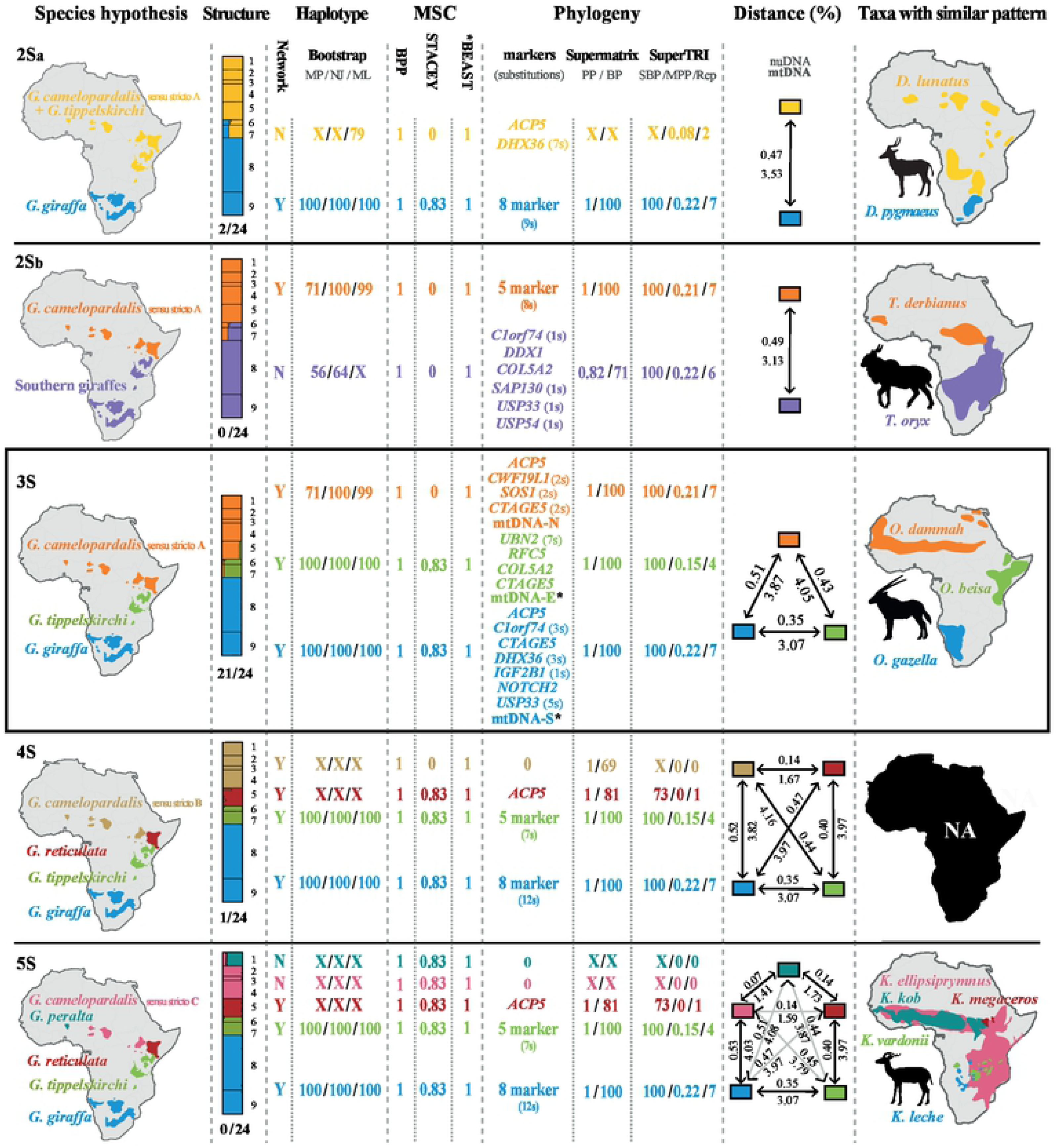
The five molecular hypotheses for giraffe taxonomy. The five taxonomic hypotheses that received some support from our analyses on giraffes show the existence of two species, with two possible geographic patterns (2Sa and 2Sb hypotheses), three species (3S hypothesis), i.e. *G. camelopardalis* sensu stricto A, *G. giraffa* and *G. tippelskirchi*, four species (4S hypothesis), i.e. *G. camelopardalis* sensu stricto B, *G. giraffa*, *G. reticulata*, and *G. tippelskirchi*, or five species (5S hypothesis), i.e. *G. camelopardalis* sensu stricto C, *G. giraffa*, *G. peralta, G. reticulata*, and *G. tippelskirchi.* In the first column are drawn the geographic distributions of giraffe species for each of the five taxonomic hypotheses. In the second column are summarized the results obtained from STRUCTURE analyses. Barplots were illustrated with DISTRUCT (1 = *peralta*, 2 = *antiquorum,* 3 = *camelopardalis*, 4 = *rothschildi*, 5 = *reticulata*, 6 = *tippelskirchi*, 7 = *thornicrofti*, 8 = *giraffa*, 9 = *angolensis*) and number of analyses supporting each taxonomic hypothesis (in total 24, see Table 3) is indicated beneath barplots. In the third column are shown the support values provided by the three Multispecies coalescent (MSC) methods, i.e. BPP, STACEY and *BEAST. In the fourth column are indicated the bootstrap values obtained with the phylogenetic analyses based on the Maximum Parsimony, Distance and Maximum Likelihood criterion (“X“: support < 50). In the fifth column are listed the markers supporting each taxonomic hypothesis in the separate analyses of 21 introns and mtDNA, as well as the support values obtained from supermatrix and SuperTRI analyses (“-“: not found). In the sixth column are detailed the mean pairwise distances between individuals of the same taxon calculated using either nuDNA data (concatenation of 21 introns, above) or mtDNA (below) (Since all the mitochondrial sequences of the subspecies *giraffa* belong to haplogroup E, they were considered as *tippelskirchi* for distance comparisons; see paragraph 4.5 for discussion on mtDNA introgression). In the seventh column are shown the distribution maps of bovid genera with a similar geographic pattern of speciation.

**Table 3.** STRUCTURE analyses based on 21 introns and associated ΔK values (highest in bold) calculated using the method of Evanno et al. [50], as well as the optimal K value(s) deduced from the “plateau” method of Pritchard et al. [44] (underlined) (Our conclusions based on the results of both methods are highlighted in grey).

| Ancestry Model | Popdata | Allele Frequency | $\Delta K$ K=2 | | $\Delta K$ K=3 | | $\Delta K$ K=4 | |
| --- | --- | --- | --- | --- | --- | --- | --- | --- |
|  |  |  | 2Sa/b hypotheses |  | 3S hypothesis |  | 4S hypothesis |  |
| | | | $\lambda = 1.0$ | $\lambda = 0.45$ | $\lambda = 1.0$ | $\lambda = 0.45$ | $\lambda = 1.0$ | $\lambda = 0.45$ |
| Admixture | - | Correlated | <u>31.5</u> <sup>a, b</sup> | 27.3 <sup>b</sup> | <u>2.2</u> | **116.5** | **49.7** | 18.3 |
| Admixture | POPID | Correlated | 28.7 <sup>b</sup> | 29.9 <sup>a, b</sup> | **1736.4** | <u>4.24</u> | 17.1 | **691.2** |
| Admixture | LOCPRIOR | Correlated | 30.6 <sup>a</sup> | 31.0 <sup>b</sup> | **614.4** | **465.9** | 10.9 | 81.3 |
| Admixture | - | independent | 24.8 <sup>a</sup> | **27.1** <sup>b</sup> | <u>4.6</u> | 4.4 | **154.9** | <u>14.7</u> |
| Admixture | POPID | independent | **25.1** <sup>b</sup> | **26.6** <sup>a</sup> | <u>4.8</u> | <u>4.5</u> | 0.8 | 5.9 |
| Admixture | LOCPRIOR | independent | **28.8** <sup>a, b</sup> | **28.8** <sup>b</sup> | <u>1.7</u> | <u>3.7</u> | <u>13.2</u> | <u>13.7</u> |
| No Admixture | - | Correlated | **12.7** <sup>a, b</sup> | <u>14.7</u> <sup>a</sup> | <u>0.3</u> | 0.3 | <u>0.3</u> | 2.3 |
| No Admixture | POPID | Correlated | 28.0 <sup>b</sup> | 26.7 <sup>a</sup> | **1556.3** | <u>4.2</u> | <u>797.8</u> | **703.9** |
| No Admixture | LOCPRIOR | Correlated | **30.8** <sup>a</sup> | <u>53.0</u> <sup>a</sup> | <u>1.3</u> | <u>1.1</u> | 4.4 | <u>3.1</u> |
| No Admixture | - | independent | 24.0 <sup>a</sup> | **25.8** <sup>a</sup> | **585.9** | 2.3 | 14 | <u>22.9</u> |
| No Admixture | POPID | independent | **25.5** <sup>b</sup> | **25.3** <sup>b</sup> | <u>4.4</u> | <u>3.0</u> | <u>5.2</u> | <u>11.7</u> |
| No Admixture | LOCPRIOR | independent | **28.5** <sup>a</sup> | **27.8** <sup>a, b</sup> | <u>1.0</u> | <u>1.2</u> | 2.6 | 2.3 |

POPID = subspecies assignment for each individual; LOCPRIOR = consideration of sampling location; “XX”: K = 2 or 3; “XX”: K = 3; “XX”: K = 4; “XX”: K = 3 or 4; “XX”: K = 2, 3 or 4; a: affiliation of the subspecies *tippelskirchi* and *thornicrofti* to *G. camelopardalis* (2Sa hypothesis in Fig. 4); b: affiliation of the subspecies *tippelskirchi* and *thornicrofti* to *G. giraffe* (2Sb hypothesis in Fig. 4).

### Nuclear and mitochondrial pairwise distances

The nuDNA-G137 dataset (16968 nt) and mtDNA-GH82 dataset (1742 nt) were used to calculate pairwise distances in PAUP* v4b10 [43] (Appendix D). For the nuclear dataset, we performed calculations considering the five taxonomic hypotheses summarized in Fig. 4. For the mtDNA dataset, we primarily performed calculations based on the three main mitochondrial haplogroups, named N, E and S depicted in Fig. 1B, but considered also the five possible hypotheses of species delimitation shown in Fig. 4.

## Results

### Phylogenetic analyses of the nuclear dataset

The 21 nuclear introns were analysed independently and in combination. The phylogenetic trees obtained from the separate analyses of the 21 independent introns are detailed in Appendix B and the Bayesian nuclear tree of the concatenated dataset (17276 nt) is depicted in Fig. 1A. The results of other analyses (ML bootstrap [BP] and SuperTRI indices [SBP/MPP/NRep]) are indicated only for the nodes supported by posterior probability (PP) values ≥ 0.9, as well as for nodes discussed in the text (e.g., subspecies).

The monophyly of *Giraffa* is supported by all analyses and almost all markers separately (NRep = 20), and the genus is diagnosed by 158 exclusive synapomorphies in the nuclear genes. Within *Giraffa*, 19 nodes are supported by PP ≥ 0.9 in the Bayesian tree of the nuDNA supermatrix (Fig. 1A; Appendix B2.22), but only three of them are associated with BP > 90: (1) the clade here named *G. camelopardalis* sensu stricto A, which groups together all members of the subspecies *camelopardalis*, *antiquorum*, *peralta*, *reticulata*, and *rothschildi* (PP = 1; BP = 100); (2) *G. giraffa*, including all members of the subspecies *angolensis* and *giraffa* (PP = 1; BP = 100); and (3) *G. tippelskirchi*, comprising all members of the subspecies *thornicrofti* and *tippelskirchi* (PP = 1; BP = 100). The monophyly of other taxa was less supported in the ML analysis: BP = 69 for *G. camelopardalis* sensu stricto B (*G. camelopardalis* s.s. A excluding *reticulata*) and BP = 81 for *G. reticulata*.

The results of separate analyses of the 21 introns showed that none of them supports the monophyly of *G. camelopardalis* s.s. B and that *G. reticulata* is found monophyletic only for *ACP5,* but with insignificant support (PP = 0.03). By contrast, *G. tippelskirchi* is independently supported by four genes: *COL5A2* (PP = 0.96), *CTAGE5* (PP = 0.64), *RFC5* (PP = 0.75) and *UBN2* (PP = 1); and all individuals of this taxon share seven molecular signatures in the *UBN2* gene (Fig. 1A). The taxa corresponding to *G. camelopardalis* s.s. A and *G. giraffa* are the most robust and reliable nodes within *Giraffa* (Fig. 1A, Appendix B2.22): *G. camelopardalis* s.s. A is supported by the separate analyses of 4 introns, i.e. *ACP5* (PP = 0.75)*, CTAGE5* (PP = 1)*, CWF19L1* (PP = 1) and *SOS1* (PP = 1), and members of this group share eight molecular signatures detected in five markers; *G. giraffa* is found monophyletic with PP ≥ 0.5 in the separate analyses of 5 introns, i.e. *C1orf74* (PP = 1), *DHX36* (PP = 0.98), *IGF2B1* (PP = 0.9), *NOTCH2* (PP = 0.5) and *USP33* (PP = 1), and members of this group share 12 molecular signatures detected in four markers.

### SuperTRI analyses

The SuperTRI analyses of the 21 introns are highly informative for relationships within *Giraffa* (Appendix B). Indeed, only six nodes are supported by MPP > 0.1 and NRep ≥ 2 (Fig. 1A): *Giraffa* + *Okapia* (MPP = 1; NRep = 21); *Giraffa* (MPP = 0.93; NRep = 20); *G. giraffa* + *G. tippelskirchi* (MPP = 0.22; NRep = 6); *G. giraffa* (MPP = 0.22; NRep = 7); *G. camelopardalis* s.s. A (MPP = 0.21; NRep = 4); and *G. tippelskirchi* (MPP = 0.15; NRep = 4). All these nodes are also characterized by several exclusive synapomorphies detailed in Fig. 1A. By contrast, SuperTRI analyses did not provide support for the two other taxa: *G. camelopardalis* s.s. B (MPP/ NRep = 0) and *G. reticulata* (MPP = 0; NRep = 1). Particularly relevant is the fact that SuperTRI results also show no support (i.e. MPP ≤ 0.05 and NRep ≤ 1; Appendix B) for all interpopulational or interindividual relationships within the three species *G. camelopardalis* s.s. A, *G. giraffa* and *G. tippelskirchi*.

### STRUCTURE analyses

Our Bayesian population structure analyses were carried out on alleles inferred for 21 introns and 137 giraffes (0.5 % of missing data). We tested different models (admixture versus no admixture, independent versus correlated allele frequency), with and without supporting priors on the subspecies (POPID) or on the geographic origins of the individuals (LOCPRIOR), as well as two values of lambda, fixed (λ = 1) or estimated (λ = 0.45) (Table 3). For each run, the most likely number of distinct groups (K) was determined using both ΔK and “plateau” methods [50,44].

Using the ΔK method of Evanno et al. [50], 58% of the STRUCTURE analyses (14 / 24) resulted in the highest ΔK value for the separation into two clusters (K) corresponding to a North/South dichotomy and the comparisons between DISTRUCT barplots indicated differences in the affiliation of both *tippelskirchi* and *thornicrofti* giraffes to either the northern or the southern group (Table 3; 2Sa and 2Sb hypotheses in Fig. 4). The highest ΔK value for three distinct clusters was obtained for 25 % (6 / 24) of the analyses (Table 3), supporting the 3S hypothesis (Fig. 4). Finally, the separation into four K clusters was supported by four analyses (17 %, Table 3).

Using the “plateau” method of [44], we found that K = 3 is the most probable number of clusters for 12 STRUCTURE HARVESTER diagrams (50% of the 24 analyses), whereas the highest support for four clusters could only be found in 8 % of the analyses (2 / 24) (Appendix C). For other diagrams, it was difficult to determine at which K the plateau is reached: for 29 % of the analyses (7 / 24), it was not possible to choose between K = 3 and 4; for 8 % of the analyses (2 / 24) it was not possible to choose between K = 2 or 3; and for 4 % of the analyses (1 / 24) it was not possible to choose between K = 2, 3 or 4.

### Analyses of nuclear haplotypes

The haplotype network and bootstrap values obtained from the ML, MP and NJ analyses of the 274 nuclear haplotypes of 137 individuals are shown in Fig. 2. All analyses support a division into three divergent haplogroups (separated by a minimum of 36 mutations) corresponding to (1) *G. camelopardalis* s.s. A (BP_MP/NJ/ML_ = 71/100/99), which includes the subspecies *camelopardalis, antiquorum, rothschildi, reticulata* and *peralta;* (2) *G. tippelskirchi* (BP_MP/NJ/ML_ = 100), which includes the subspecies *tippelskirchi* and *thornicrofti*, and (3) *G. giraffa* (BP_MP/NJ/ML_ = 100) containing the southern subspecies *giraffa* and *angolensis*.

The haplotype network shows a separation between *reticulata* and other subspecies of *G. camelopardalis* s.s. A (a taxon named *G. camelopardalis* s.s. B in Fig. 4), as well as a separation between the two subspecies of *G. tippelskirchi*, i.e. *tippelskirchi and thornicrofti*. None of these additional clusters are however supported by BP_MP/NJ/ML_ > 50, except *G. camelopardalis* s.s. B (BP_ML_ = 72) and *thornicrofti* (BP_ML_ = 54) in the RAxML analysis. By contrast, no subspecies can be distinguished within *G. giraffa*.

The haplotype networks constructed for each of the 21 nuclear introns are shown in Fig. 3. Only five taxa show allelic clustering: (1) *G. camelopardalis* s.s. A and (2) the group *G. giraffa* + *G. tippelskirchi* in seven networks (*C1orf74*, *CTAGE5*, *CWF19L1*, *SAP130*, *SOS1*, *USP33* and *USP54*); (3) *G. giraffa* in six networks (*ACP5*, *C1orf74*, *DHX36*, *IGF2B1*, *RFC5*, and *USP33*); (4) *G. tippelskirchi* in six networks (*C1orf74*, *COL5A2*, *CTAGE5*, *RFC5*, *UBN2*, and *USP33*); and (5) *thornicrofti* in one network (*IGF2B1*).

We detected incomplete clustering (i.e., 1-3 “foreign” alleles in the cluster, or less than three alleles not included into the cluster) for the following taxa: *G. camelopardalis* s.s. A (*ACP5*: one *thornicrofti* allele; *DDX1* and *RFC5*: two alleles outside); *G. giraffa* (*DDX1*: two *reticulata* alleles; *NOTCH2*: one *tippelskirchi* allele; *SOS1*: two alleles outside; *USP54*: two *thornicrofti* alleles); *G. tippelskirchi* (*ACP5*: one allele outside; *SOS1*: two *giraffa* alleles); *G. camelopardalis* s.s. B (*USP54*: three *reticulata* alleles); and *G. reticulata* (*ACP5* and *USP54*: three alleles outside).

The patterns found for the six other introns (*CCT2*, *MACF1*, *NUP155*, *OTOF*, *PLCE1*, *RASSF4*) do not fit any taxon depicted in the hypotheses of Fig. 4.

### Multispecies coalescent analyses

We constructed a MSC species-tree from the nuDNA-G274-O6 dataset using *BEAST. The topology is similar to the supermatrix topology of Fig. 1A, with maximal support (PP = 1) for *G. camelopardalis* s.s. A, *G. giraffa* and *G. tippelskirchi.* However, the monophyly of *G. camelopardalis* s.s. B, *G. camelopardalis* s.s. C and four subspecies (*antiquorum*, *peralta, reticulata*, and *thornicrofti*) was also highly supported (PP = 1) in the MSC tree (Appendix E2). The subspecies *tippeskirchi* was found monophyletic, but with low PP support (= 0.39).

The analyses based on STACEY showed highest support for five distinct giraffe species, i.e., *G. camelopardalis* s.s. C*, G. giraffa, G. peralta, G. reticulata* and *G. tippeskirchi*, a pattern found in 87 % of the trees. Other hypotheses of species delimitation were less supported: the 4S hypothesis (*G. camelopardalis* s.s. B*, G. giraffa, G. reticulata* and. *G. tippeskirchi*) was found in 7% of the trees; whereas the 6S hypothesis, which recognizes *G. camelopardalis* s.s. C, *G. giraffa, G. peralta, G. reticulata, G. tippeskirchi* sensu stricto, and *G. thornicrofti*, was found in 6% of the trees. Similar results were obtained when outgroup sequences were excluded (data not shown).

Species delimitation analyses based on BPP provided maximal support (PP = 1) for all species recognized according to the 3S, 4S, and 5S hypotheses (Fig. 4). The same results were found with the three species model priors (SMP1, 2 or 3; Appendix E1). The further division of *G. tippelskirchi* into two separate taxa, i.e. *G. tippelskirchi* sensu stricto and *G. thornicrofti* (6S hypothesis) was only weakly supported (PP_SMP1_ = 0.26; PP_SMP2_ = 0.4; PP_SMP3_ = 0.34).

### Phylogenetic analyses of the mitochondrial fragment

The Bayesian tree reconstructed from the mtDNA-GH82O3 dataset (1776 nt) is shown in Fig. 1B. It shows the existence of three major geographic haplogroups: northern (N), eastern + southeastern (E), and southwestern (S) giraffes.

The N haplogroup is supported by both Bayesian and bootstrap analyses (PP = 1; BP = 93). It includes all haplotypes detected for *G. camelopardalis* s.s. A, as well as one divergent haplotype of *G. tippelskirchi* (TIP15, EU088334) sequenced by Brown et al. [29] for nine individuals from Kenya (Athi River Ranch) (see details in Appendix B1.1). Three subspecies of *G. camelopardalis* are monophyletic: *antiquorum* (PP = 1; BP = 85), *peralta* (PP = 1; BP = 99) and *rothschildi* (PP = 1; BP = 83). The subspecies *camelopardalis* is found polyphyletic. The reticulated giraffes constitute a polyphyletic assemblage: although most of them are grouped together (PP = 1; BP = 92) as the sister group of the divergent haplotype TIP15 (EU088334) of *G. tippelskirchi* (PP = 1; BP = 94), the haplotype RET8 sequenced by Fennessy et al. [38] is closely related to *rothschildi* (PP = 0.89; BP = 46), and the haplotype RET9 (EU088321) sequenced by Brown et al. [29] appears as the sister group of all other northern haplotypes.

The E haplogroup comprises giraffes from eastern and southeastern Africa (PP = 0.99; BP = 87). It contains members of two putative species, *G. tippelskirchi* and *G. giraffa*, and can be further divided into three subgroups corresponding to “Masai I”, “Masai II”, and the subspecies *giraffa.* The interrelationships between the three subgroups are unresolved. The Masai I subgroup (PP = 1; BP = 95) contains Masai giraffes (subspecies *tippelskirchi*) from Kenya and Tanzania. The Masai II subgroup (PP = 1; BP = 89) includes Masai giraffes (subspecies *tippelskirchi*) from Kenya and Tanzania, as well as giraffes of the subspecies *thornicrofti* from northern Zambia (Luangwa Valley National Park). The third subgroup represents the subspecies *giraffa* (PP = 1; BP = 99) and includes giraffes from southern Zambia, northern Botswana, northeastern Namibia, Zimbabwe and South Africa.

The S haplogroup contains exclusively individuals of the subspecies *angolensis* from Namibia and central Botswana. Its monophyly is less supported than the two other mitochondrial haplogroups (PP = 0.37; BP = 60). Our analyses provide a moderate support (PP = 0.94; BP = 65) for an early divergence of the S haplogroup.

### Nuclear and mitochondrial pairwise distances

The alignment of 21 nuclear introns was used to calculate pairwise distances between giraffes (Fig. 4 and Appendix D2). The results show that the mean distance between *G. camelopardalis* s.s. B and *G. reticulata* is 0.14 % and the mean distance between *G. camelopardalis* s.s. C and *G. peralta* is 0.07 %, which is significantly smaller than other interspecific distances involving *G. camelopardalis* s.s. A, *G. giraffa* and *G. tippelskirchi* (comprised between 0.35 and 0.51 %).

For the mtDNA alignment, we calculated pairwise distances between 82 haplotypes. Three haplotypes (TIP15, RET8 and RET9) were excluded from the analysis due to their grouping outside of their assigned taxon in the phylogenetic tree (Fig. 1B). The distances between the haplogroups identified in Fig. 1B are summarized in Appendix D1 and Fig. 4. There are three major haplogroups: haplogroup N= northern (= *G. camelopardalis* s.s. A); haplogroup E = Masai I, Masai II, and southeastern (= subspecies *giraffa*); and haplogroup S= southwestern (= subspecies *angolensis*). The mean distances between these three haplogroups are comprised between 3.07 and 4.16 %. Within haplogroup N, the distances between *G. camelopardalis* s.s. B and *reticulata* range from 1.29 % (ROTH3 *versus* RET3) to 2.19 % (PER2 *versus* RET13). Within haplogroup E, we found similar distances between Masai I, Masai II and southeastern haplotypes, i.e., between 1.17 % (TIP1 *versus* GFA7) and 2.12 % (TIP5 *versus* GFA9). Within haplogroup S, the distances range from 0 to 0.96 % (ANG12 *versus* ANG16).

## Discussion

### Population genetic analyses support the 3S hypothesis

The assessment of population genetic structure has become indispensable in evolutionary biology and conservation to reveal hidden biodiversity. Among freely accessible software provided for this task, STRUCTURE [26] is the most commonly used program, with 17473 citations in Web of Science (January 2019). Using Bayesian inference, STRUCTURE is a model-based clustering method to detect population structure and assign individuals to K populations [26]. However, many published results based on STRUCTURE are not reproducible because the genotypes were not available or the parameters used for the analyses were not fully detailed by the authors [45,61].

The program STRUCTURE was previously used to infer genetic structure in giraffe populations, using either genotypes from 14 microsatellite loci of 381 individuals [29] or PHASED alleles of seven introns for 105 giraffes [38] or rather the extended dataset of 21 introns for 137 individuals [12]. Brown et al. [29] suggested the existence of at least six species, but the optimal K was not determined using either the method of Evanno et al. [50] or that of Pritchard et al. [44], and their results are not reproducible, because the microsatellite data were not made available. According to Winter et al. [12], “K = 4 shows four well resolved groups and is supported as best fitting number of clusters by several statistical methods”, but they did not provide any details on the model and method used for their STRUCTURE analyses. Using the same dataset, comprising allelic information of 21 nuclear introns for 137 giraffes, we tested 16 different models under STRUCTURE in order to shed more light on giraffe population structure. Considering the method of Evanno et al. [50], 58% of the analyses provided support for two distinct populations of origin (K = 2), 25% for three distinct clusters (K = 3), and only 17% confirmed the result obtained by Winter et al. [12], i.e. K = 4.

The selection of the appropriate K using the method of Pritchard et al. [44] partly confirmed previously mentioned difficulties to determine the point of plateau [46,50]. We clearly recognize K = 3 as the optimal clustering for 50% of the analyses. For other analyses, it was difficult to identify at which K the plateau is reached (K = 2 or 3?; K = 2, 3 or 4?; K = 3 or 4?; K = 3, 4 or 5?; Appendix C; Table 3).

Selecting the best suitable model for STRUCTURE is far from simple, especially for taxa with a wide distribution range like giraffes. The choice of an admixture model with correlated allele frequency seems appropriate for populations of East Africa, where hybrids between individuals from divergent populations were previously described (see below). However, such a model may be more questionable for isolated populations, such as the subspecies *peralta*. In order to better estimate the optimal value of K under STRUCTURE, we recommend therefore for future users of the program to test different combinations of model parameters, to estimate the value of λ, and to make comparison between optimal K estimated with either the ΔK method [50] or the “plateau” method [44]. Using this approach and taking into account that the ΔK method can be biased towards K = 2 [61] and that the smallest value of K is preferred when several values of K give similar estimates of log Pr (X | K) [44], we concluded that K = 3 is the most likely hypothesis for 88 % of the analyses (highlighted in grey in Table 3).

Our network and bootstrap analyses of the 274 nuclear giraffe haplotypes (21 introns, 137 giraffes), as well as the networks of the 21 introns, also highly support a division into three divergent haplogroups, representing the three species *G. camelopardalis* s.s. A, *G. giraffa*, and *G. tippelskirchi* (Fig. 2 and 3).

### Phylogenetic analyses support the 3S hypothesis

In the nuclear tree reconstructed from the concatenation of 21 introns (Fig. 2), four putative species were found to be monophyletic: *G. giraffa*, *G. tippelskirchi*, *G. camelopardalis* s.s. A and *G. reticulata.* However, the two latter mentioned taxa obtained weak ML bootstrap support (BP = 69 and 81, respectively). To further investigate phylogenetic relationships, we conducted separate Bayesian analyses for all markers and summarized the results with the SuperTRI method [20]. Within *Giraffa*, the analyses showed that only four nodes can be considered as reliable (SBP = 100; MPP > 0.15; Nrep > 4): *G. camelopardalis* s.s. A (grouping together northern and reticulated giraffes), *G. giraffa* (southern giraffes), *G. tippelskirchi* (southeastern giraffes), and *G. giraffa* + *G. tippelskirchi* (Fig. 2). All these nodes are supported by the separate analyses of several independent introns (between four and seven), which explain why MPP values are significantly higher than for all intraspecific relationships (between 0.15 and 0.22 versus between 0 and 0.03). By contrast, the SuperTRI analyses provided no support (MPP = 0; Nrep ≤ 1) for the existence of both *G. camelopardalis* s.s. B and *G. reticulata*. The monophyly of *G. reticulata* was found by only *ACP5,* but with insignificant support (PP = 0.03).

### Multispecies coalescent approaches show further geographic structure

Two MSC methods, *BEAST and BPP, showed strong support (PP = 1) for the 3S hypothesis, in which three species can be distinguished, i.e., *G. camelopardalis* s.s. A, *G. giraffa*, and *G. tippelskirchi.* However, STACEY analyses provided support for further species delimitation, i.e., the 5S hypothesis (87%). The five taxa, *G. camelopardalis* s.s. C, *G. giraffa*, *G. peralta*, *G. reticulata*, and *G. tippelskirchi*, are also highly supported by both *BEAST and BPP analyses (PP = 1). As recently pointed by Sukumarana and Knowles [62] and Jackson et al. [63], it appears that multispecies coalescent methods delimit structure, not species. In agreement with that, it is important to note that only two of the five putative MSC species can be diagnosed by molecular signatures (Fig. 1), i.e. the ones assumed by the 3S hypothesis: *G. tippelskirchi* is characterised by seven exclusive synapomorphies (ES), all found in the *UBN2* gene, which are shared by 19 individuals; and *G. giraffa* is characterised by 12 ES detected in four independent genes and shared by 61 individuals. For the three other taxa of the MSC 5S hypothesis, we did not detect any fixed mutation in the 21 nuclear introns. This means that the populations of *G. camelopardalis* s.s. C, *G. peralta*, and *G. reticulata* have never been completely isolated genetically. Their grouping into *G. camelopardalis* s.s. A is however supported by eight ES detected in five independent genes and shared by 57 individuals. The 3S hypothesis is therefore strengthened by the criterion of genetic isolation, as the detection of ES in the three species *G. camelopardalis* s.s. A, *G. giraffa*, and *G. tippelskirchi* indicates that their populations were reproductively isolated during enough time, allowing for the fixation of diagnostic mutations in all individuals.

### Interspecies relationships within *Giraffa*

According to the fossil record, contemporary giraffes first appeared during the Pleistocene around 1 Mya [64], a hypothesis also supported by molecular dating estimates [65]. All candidate species to root the tree of giraffes are highly distant taxa: *Okapia*, which is the only other extant genus of the family Giraffidae, separated from *Giraffa* during the Middle Miocene (around 15.2 Mya); other ruminant families, such as Bovidae, Cervidae, Moschidae and Antilocapridae, diverged from Giraffidae at the transition between Oligocene and Miocene (around 23.4 Mya) [65]. The rooting of the giraffe tree can be therefore misleading due to a long branch attraction (LBA) artefact (for a review see Bergsten [66]) between the distant outgroup and one of the longest branches of the ingroup. This problem explains the highly variable root position in our mitochondrial analyses: with MrBayes, the first haplogroup to diverge is either S (Fig. 1B, PP = 0.37; BP = 60) or E (if the two bovid species are excluded as outgroup taxa, data not shown, PP = 0.55); with BEAST, haplogroups E and S are found to be sister-groups (PP = 0.74), as in the mitochondrial tree of Fennessy et al. [38].

The nuclear dataset provided more signal for resolving basal relationships within *Giraffa*. As indicated in Fig. 1, our phylogenetic analyses supported a sister-group relationship between *G. giraffa* and *G. tippelskirchi* (PP = 0.82; BP = 71). This node was found monophyletic with 6 independent markers (*C1orf74, DDX1, COL5A2, SAP130, USP33* and *USP54*). By comparison, SuperTRI analyses clearly showed that the two other hypotheses (either *G. camelopardalis* s.s. A + *G. giraffa* or *G. camelopardalis* s.s. A + *G. tippelskirchi*) are less supported (MPP ≤ 0.09; NRep ≤ 2 markers). All these results agree therefore with a deep North/ South dichotomy within *Giraffa*.

### Evidence for introgressive hybridization between giraffe species

The comparison between the mtDNA tree based on 82 giraffe haplotypes and the nuclear tree reconstructed from 21 introns sequenced for 137 giraffes reveals a robust conflict for the evolutionary history drawn from maternal and biparental markers (Fig. 1). Some mito-nuclear conflicts can be simply explained by recent hybridization between sympatric or parapatric taxa (species or subspecies), resulting in the transfer of the mitochondrial genome from one taxa to the other, a process referred to as mitochondrial introgression [6–11].

A first case of potential hybridisation is represented by the mitochondrial haplotype TIP15, which constitutes the sister-group of the main haplogroup of reticulated giraffes (Fig. 2B), from which it differs by a distance of only 1%. The nine Masai giraffes possessing this haplotype were collected in southern Kenya (Athi River Ranch) [29], where wild populations of *tippelskirchi* and *reticulata* can sometimes hybridize [67]. We suggest therefore that introgressive hybridization can account for the transfer of the mitochondrial haplotype TIP15 from *reticulata* to *tippelskirchi*. The allelic networks of the 21 nuclear introns suggest also past nuclear introgression, this time from *tippelskirchi* to *reticulata*, as two individuals of *reticulata*, ISC04 and RETWil2, are characterized by several rare alleles identical or similar to those found in *tippelskirchi*: in *ACP5* (only for RETWil2), *COL5A2* (only for ISC04), *CTAGE5* and *DDX1* (both individuals) (Fig. 3).

The second case of mitochondrial introgression concerns the haplotype RET8 detected in one reticulated giraffe from the Nürnberg Zoo [38]. Its grouping with Rothschild’s giraffes may be explained by interbreeding between *reticulata* and *rothschildi* either in zoos [68] or in the wild, as field observations have documented the occurrence of *reticulata* X *rothschildi* hybrid phenotypes in Kenya [69]. Unfortunately, these hybrid individuals or populations were not yet studied for nuclear genes.

The mitochondrial haplotype RET9, which was detected by Brown et al. [29] in a single reticulated giraffe (accession number: EU08821), is intriguing because it is divergent from all other sequences of haplogroup N. We propose two hypotheses to explain its divergence. The first hypothesis assumes the retention of ancestral haplotypes in wild populations of reticulated giraffes; it will be confirmed if identical or similar haplotypes are discovered in other reticulated giraffes. Another hypothesis implies that the sequence EU088321 is problematic, either because it contains multiple sequencing errors or because it is a nuclear sequence of mitochondrial origin (Numt) [70]. Obviously, further investigations are needed to solve this issue.

The most important and interesting mito-nuclear discordance concerns giraffes from eastern and southern Africa. In the nuclear tree (Fig. 2A), these giraffes are divided into two geographic groups corresponding to two different species: giraffes from southern Africa (South Africa, Namibia, Botswana, and southern Zambia) belong to *G. giraffa*, whereas eastern giraffes (southern Kenya, Tanzania, and northern Zambia) belong to *G. tippelskirchi*. These two species are not monophyletic in the mitochondrial tree: *G. giraffa* is polyphyletic, because members of the two subspecies *giraffa* and *angolensis* are not grouped together; whereas *G. tippelskirchi* is paraphyletic, due to the inclusive position of the subspecies *giraffa* (southeastern giraffes). To interpret these conflicting results, it is crucial to remember that basal relationships within *Giraffa* are not reliable in the mitochondrial tree, due to a high genetic distance towards outgroup taxa (see above for explanations). Taken this in mind, it can be hypothesized that the three species identified with nuclear data were characterized by three different ancestral mitochondrial haplogroups: N for *G. camelopardalis* s.s. A, E for *G. tippelskirchi*, and S for *G. giraffa*. According to this hypothesis, we can further propose that the common ancestor of southeastern populations of *G. giraffa* (subspecies *G. g. giraffa*) acquired a mitochondrial genome from *G. tippelskirchi* (haplogroup E) by introgressive hybridization between parapatric populations. Using a calibration at 1 ± 0.1 Mya for the common ancestor of giraffes [64–65], we estimated that the introgressive event occurred around 420 kya (see Appendix B1.4), i.e. during one of the most important glacial periods of the Pleistocene. In sub-Saharan Africa, glacial periods were generally characterized by the contraction of forest areas and the concomitant extension of open areas, such as savannahs and deserts. In addition, river levels were lower, facilitating dispersals and the colonization of new areas. Since Pleistocene environments were more stable in subtropical southern East Africa than in tropical East Africa [71], we suggest that some Masai giraffes migrated around 420 kya from East Africa to southern East Africa, promoting secondary contacts between *G. tippelskirchi* and *G. giraffa*, and therefore the mitochondrial introgression of haplotype E into. *G. g. giraffa*. In the latter subspecies, the ancestral haplotype S has been completely replaced by the new haplotype E. By contrast, the ancestral haplotype S has been maintained in southwestern populations of the subspecies *G. g. angolensis*. The absence of haplotype E in southwestern giraffes suggests that female giraffes were not able to disperse from East to West and reciprocally. Important biogeographical barriers may have been the Kalahari Desert during glacial periods of the Pleistocene, and the Okavango Delta associated with Palaeo-lake Makgadikgadi during interglacial periods. However, nuclear data support gene flow mediated by dispersing males between eastern (*G. g. giraffa*) and western populations (*G. g. angolensis*) of southern giraffes. Female philopatry and male biased dispersal are classically observed in mammal species [72]. In giraffes, such different sexual behaviours can be explained by nursery herds, which consist of several females and their offspring [73], and by solitary males, which spend a lot of time to find receptive females. Thereby, males may often have to migrate over long distances to successfully pass on their genes [74]. In this regard, we can assume that males are generally more willing than females to take the risk of overcoming biogeographic barriers, such as deep rivers or large deserts. Markers from the Y chromosome should be sequenced to further address our biogeographic scenario involving a better dispersal capacity for males than females.

### Conclusion for giraffe conservation management

The species is the most important taxonomic unit for conservation assessments and for the establishment of justified management plans [75–76]. Giraffes are currently considered as a single species by the IUCN [37], but its status has recently moved from Least Concern to Vulnerable due to a population decline of 36-40% over three generations. Even though, the situation seems to have improved for some populations (e.g. *giraffa* [77]; *peralta* [78]) in the course of enhanced conservation management, population numbers of most subspecies continue to decrease [37].

Our taxonomic study indicates that the conservation status should be separately assessed for the three species *G. camelopardalis* s.s. A (northern giraffes), *G. giraffa* (southern giraffes) and *G. tippelskirchi* (Masai giraffes). According to population estimations of the IUCN [37], the southern species *G. giraffa,* has recently increased by 168% and hence fall into the category “Least Concern”; the East African species *G. tippelskirchi* has decreased by ≥ 50% over a period of three generations and hence should be listed as “Vulnerable”; the northern species *G. camelopadalis* s.s. A has decreased by ≥70 % over the past 30 years and with only 20 000 individuals left in the wild, it should be listed under the category “Endangered” (according to Criterion A1 [79]).

## Acknowledgments

AH would like to thank Klaus-Peter Koepfli and Robert K. Wayne, who provided additional information and clarification on their sequences published in 2007. The PhD thesis of AP was funded by the LabEx BCDiv.

## Supplementary material

Supplementary data associated with this article can be found, in the online version, at https://

Appendix A. Classifications of *Giraffa*

Appendix B. Phylogenetic analyses

Appendix C. Population Structure analyses

Appendix D. Genetic distances

Appendix E. Multispecies coalescent approach

Appendix F. Analyses of nuclear haplotypes

